# Fractionated follow-up chemotherapy delays the onset of resistance in bone metastatic prostate cancer

**DOI:** 10.1101/274704

**Authors:** Pranav I. Warman, Artem Kaznatcheev, Arturo Araujo, Conor C. Lynch, David Basanta

**Affiliations:** Duke University, Durham, NC, USA; Integrated Mathematical Oncology, H. Lee Moffitt Cancer Center and Research Institute, Tampa, FL, USA; Department of Computer Science, University of Oxford, Oxford, UK; Department of Translational Hematology & Oncology Research, Cleveland Clinic, Cleveland, OH, USA; Department of Tumor Biology, H. Lee Moffitt Cancer Center and Research Institute, Tampa, FL, USA

**Keywords:** evolutionary game theory, prostate cancer bone metastasis, chemotherapy, heterogeneity, resistance

## Abstract

Prostate cancer to bone metastases are almost always lethal. This results from the ability of metastatic prostate cancer cells to co-opt bone remodeling leading to what is known as the *vicious cycle*. Understanding how tumor cells can disrupt bone homeostasis through their interactions with the stroma and how metastatic tumors respond to treatment is key to the development of new treatments for what remains an incurable disease. Here we describe an evolutionary game theoretical model of both the homeostatic bone remodeling and its co-option by prostate cancer metastases. This model extends past the evolutionary aspects typically considered in game theoretical models by also including ecological factors such as the physical microenvironment of the bone. Our model recapitulates the current paradigm of the *”*vicious cycle*”* driving tumor growth and sheds light on the interactions of heterogeneous tumor cells with the bone microenvironment and treatment response. Our results show that resistant populations naturally become dominant in the metastases under conventional cytotoxic treatment and that novel schedules could be used to better control the tumor and the associated bone disease compared to the current standard of care. Specifically, we introduce fractionated follow up therapy – chemotherapy where dosage is administered initially in one solid block followed by alternating smaller doeses and holidays – and argue that it is better than either a continuous application or a periodic one. Furthermore, we also show that different regimens of chemotherapy can lead to different amounts of pathological bone that are known to correlate with poor quality of life for bone metastatic prostate cancer patients.

## Introduction

Prostate cancer (PCa) is the second most common type of cancer in men, with over 160,000 men diagnosed during 2017 in the US alone, of which over 26,000 succumbed[1]. The majority of patients that die of the disease does so because of distant metastases, 90% of which are to the bone. For this reason, a better understanding of tumor-bone interactions is key if we are to improve how bone metastatic prostate cancer patients are treated.

Improvements in our understanding of the molecular mechanisms involved in this disease have led to the discovery of new therapeutic targets, but it remains lethal and resistance emerges in virtually every patient. Thus understanding the impact of treatments in a complex heterogeneous tumor requires approaches that can incorporate several scales of biological insights. A successful approach needs to recapitulate the emergent process of metastatic prostate cancer establishment and the emergence of resistance to treatment. In the past we demonstrated how a sophisticated agent-based computational model could help us understand the role of PCa-host interactions, as well as the impact of existing and novel treatments [2, 3]. While agent-based models can accurately and quantitatively recapitulate cell dynamics in a specific area of a tumor, capturing the relevant intra-tumor heterogeneity and sensitivity to treatments can be a challenging and time-consuming process [4]. Alternatively, non-spatial population models that capture intra-tumor heterogeneity can be used to represent the entire metastatic burden of the disease and thus could be used as proxies for the patient in optimization algorithms [5]. Further, non-spatial models lend themselves to easier measurement and operationalization [6], and avoid the confusion of population-level intuitions for reductive ground truth [7].

Simple qualitative models can be useful in providing an understanding of how certain interactions can shape evolution and resistance in cancer. Previously, Ryser et al. used a simple ordinary differential equations (ODEs) model to understand bone remodeling [8, 9], and to illustrate how tumor cell interactions with bone remodeling cells can shape cancer progression [10]. Evolutionary game theory (EGT) is a particularly powerful, yet simple and qualitative, approch to focusing on the role of interactions in cancer. It originated in looking at the effects of heterogeneity [11, 12] and we have used the it to look at a wide number of dynamics in cancer like go-vs-grow [13, 14], Warburg effect [15], tumour-stroma interaction [16], and interaction of multiple public goods [17]. In the process, we have built EGT models of many cancers, including prostate cancer [15, 16] – as have several other groups. Dingli et al. [18] focused on a type of bone cancer, multiple myeloma, using an EGT approach that incorporated both tumor cells as well as bone stroma cells. More recently, West et al. [19] studied the impact of novel treatments like adaptive therapies in the context of bone metastatic prostate cancer but without including the cellular species that characterize the bone ecosystem.

In this work, we aim to model not only cancerous growth in the bone but capture key aspects of normal bone homeostasis and its co-option. We built an EGT model where PCa cells can co-opt cells from the Bone Modeling Unit (BMU). In brief, the BMU is an area of trabecular bone that is being remodeled via osteoclasts (OCs) and osteoblasts (OBs), which resorb and deposit bone tissue respectively, and the interplay between these and other BMU cell types via molecules such as receptor activator of nuclear-factor kappa-B ligand (RANK-L), which is generated by osteoblasts and promotes osteoclast differentiation and survival, and TGF-*β*, which is released when bone tissue is removed by Osteoclasts. These cellular species make the bone a very dynamic organ [20]. Furthermore, studies have shown that metastatic cancer cells can co-opt this process for their own benefit [21].

Capturing this aspect of regulation and dis-regulation of bone homeostasis allows us to better understand the selection that drives metastatic cancer evolutionary dynamics and the impact of treatments on the bone microenvironment – a topic. where the first summand of *W*OB corresponds to the ben-efit conferred to OCs by OBs that can be attributed to the secretion of RANKL by OBs that results in the recruitment of OCs to the remodeling site. This is proportional to the proportion of OCs among healthy cells (*p*OC = *ρ*OC) and lack of bone 1 *- B*. In the second summand, *δ* is the benefit that an OB receives from interacting with any PCa cell as the PCa cells secrete modest amounts of TGF-*β*.which captures the benefit conferred to OB by OC that can be attributed to the secretion of TGF-*β* as a result of bone resoption. This is proportional to the proportion of clinical significance. Furthermore, we also incorporate ecological aspects by modeling the role of the bone in regulating the fitness of the different cell types [22]. Our results suggest that we can optimize the combination of treatment on and off periods to limit tumor growth and control bone growth.

## Model

The interactions between osteoclasts (OC) and osteoblasts (OB) orchestrate bone remodeling and homeostasis. As such, bone volume is determined by the balance between the density of OBs (*ρ*OB) and density of OCs (*ρ*OC) on the existing bone, where OBs increase the bone volume and OCs decrease the bone volume according to the following discrete time dynamics:

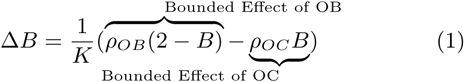

where OBs and OCs are in the range *B*(*t*) *∈* [0, 2] and *K* is a constant for determining time-scales.

Note that the above dynamics are physical and not evolutionary. The dynamics of densities for OB, OC, and the sensitive and resistant tumours (*ρT, ρTR*) however are ecological and given by logistic growth:

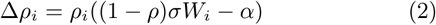

where *i ∈ {*OB, OC, *T, TR}* and *ρ* = Σ*i ρi* is the total density (thus 1 *- ρ* is the amount of remaining space in the niche), *σ* is the selection strength, *α* is the death rate, and *Wi* are the fitness functions

- The fitness function for OB is given by:

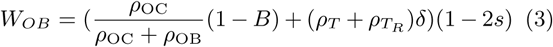

where the first summand of WOB corresponds to the benefit conferred to OCs by OBs that can be attributed to the secretion of RANKL by OBs that results in the recruitment of OCs to the remodeling site. This is proportional to the proportion of OCs among healthy cells 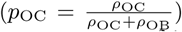 and lack of bone 1 – B. In the second summand, is the benefit that an OB receives from interacting with any PCa cell as the PCa cells secrete modest amounts of TGF-β
- The fitness function for OC is given by:

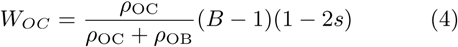

which captures the benefit conferred to OB by OC that can be attributed to the secretion of TGF β as a result of bone resoption. This is proportional to the proportion of OBs among healthy cells 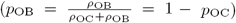 and overabundance of bone B–1. Since both WOB and WOC are functions of proportions of OC and OB, we could rewrite their coupling as a replicator dynamic. See Kaznatcheev [23] for an example of this in a similar model.
- Finally, the fitness functions for the chemotherapy sensitive and resistant tumour are:

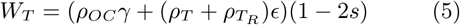

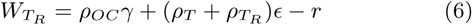

where *γ* is the benefit a PCa cell receives from interacting with OCs. The OC-led resorption of the bone allows the neighboring PCa cells to access nutrients and growth factors previously embedded in the bone; *E* is the benefit that a PCa cell receives from interacting with other PCa cells derived from the secretion of TGF-*β*. The cost of resistance to treatment is *r*. Resistance to chemotherapy is common and results from the treatment providing strong selection for PCa cells that can avoid its cytotoxicity. This resistance often comes through the up-regulation of drug exporter pumps on the surface of PCa cells. Producing and maintaining these pumps is energetically costly to the PCa cells but allows those that have a sufficient number of them to deal with cytotoxic drugs.

### Chemotherapy

For all the fitness functions above, chemotherapy is implemented by the introduction of a cost *s*, which corresponds to the the strength of the chemotherapy regiment. Chemotherapy is widely used in treating hormone-sensitive and insensitive metastatic prostate cancer [24]. As chemotherapy options like docetaxel become more widely used, so does the importance of understanding how to better administer them so as to minimize the possibility that resistant phenotypes emerge. As a simple caricature, it can be described microdynamically as: when a cell tries to undergo mitosis, it will be killed with probability *s*. This means that at *s* = 0.5, there will be no growth in the affected population, since half the time that a cell tries to divide, it succeeds and becomes two cells and half the time it dies and becomes zero cells. For 0 *< s <* 0.5, the growth rate is slowed; and for 0.5 *< s <* 1 the growth rate is reversed. This is implemented in all fitness functions (except for chemo-resistant *TR*) by multiplying fitness by (1 *-* 2*s*). Note that chemotherapy only impacts the bone via interfering with the cells that control bone perturbation, i.e. OB and OC.

## Results

We will illustrate our results with simulations of a particular parameter settings given by death rate *α* = 0.005, selection strength *σ* = 0.05, bone adjustment rate 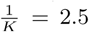, and tumour-environment ecological interactions of *δ* = 1.5, *γ* = 300, *ϵ* = 0.03.

With the model described we can recapitulate bone homeostasis when there are no PCa cells. Assuming initial densities of the different population as follows: *ρ*OB(0) = 0.001, *ρ*OC(0) = 0.01, *ρT* (0) = *ρTR* (0) = 0.0, *B*(0) = 1. The plots shown in figure 1 show how the bone is initially resorbted, and then deposited as OCs and OBs work to restore the balance of bone to homeostatic levels after a simulated microfracture.

**Figure 1:**
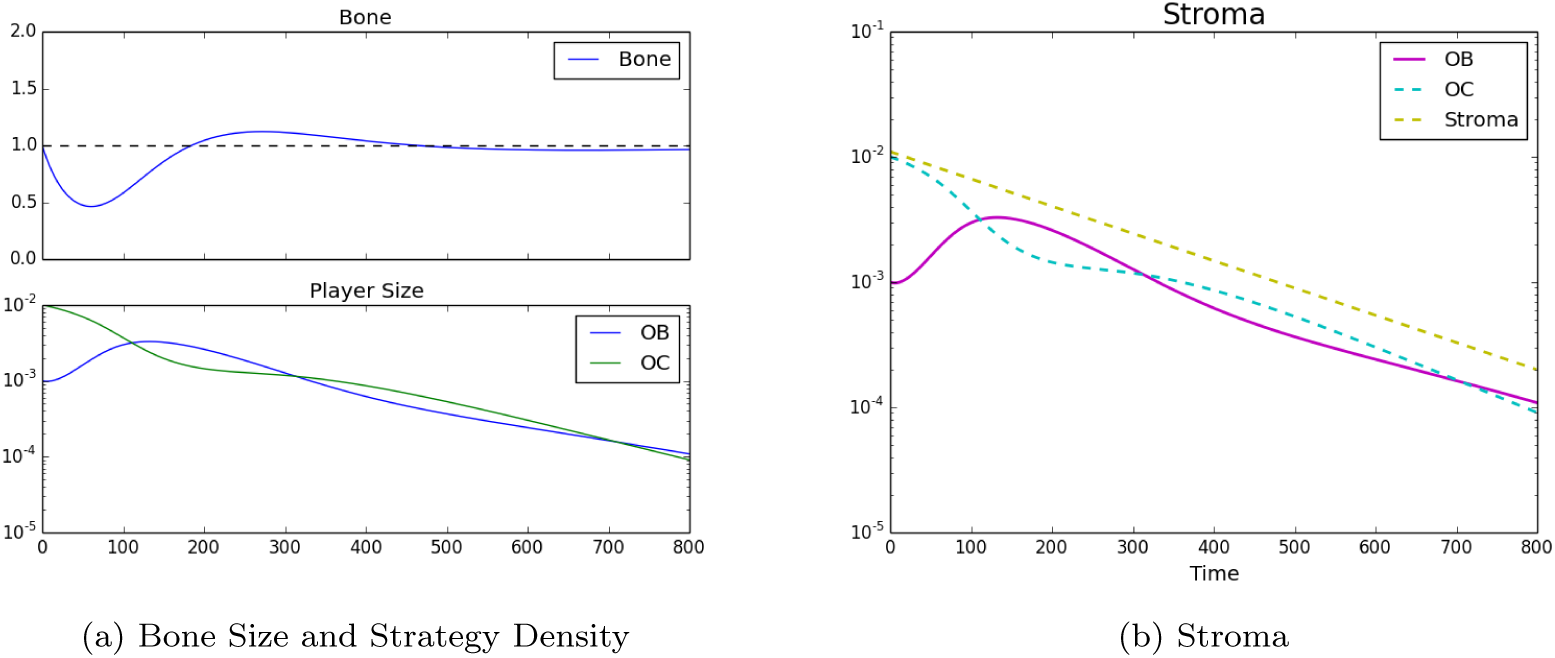
Homeostatic populations were observed by setting strategies to *ρ*OB(0) = 0.001, *ρ*OC(0) = 0.01, *ρT* (0) = *ρTR* (0) = 0.0, *B*(0) = 1. Furthermore, in subfigure (a) the decrease in stroma after the bone remodeling event. In subfigure (b) we see the classic bone fluctuation characteristic of bone remodeling units and no distinct strategies leftover, note the log-scale for the bottom subplot.

### Tumor Introduction

We now assume that a metastatic PCa cell has extravasated into an area of the bone that will undergo remodeling. In this case we assume that the initial conditions are: *ρ*OB = 0.001, *ρ*OC = 0.01, *ρT* = 0.0005, *ρTR* = 0.0001, *B*(0) = 1. In this case the results can be seen in figure 2, which shows how the growth of the PCa population leads to an increase of pathological bone and a dominance of the PCa cells over the OBs and the OCs.

**Figure 2:**
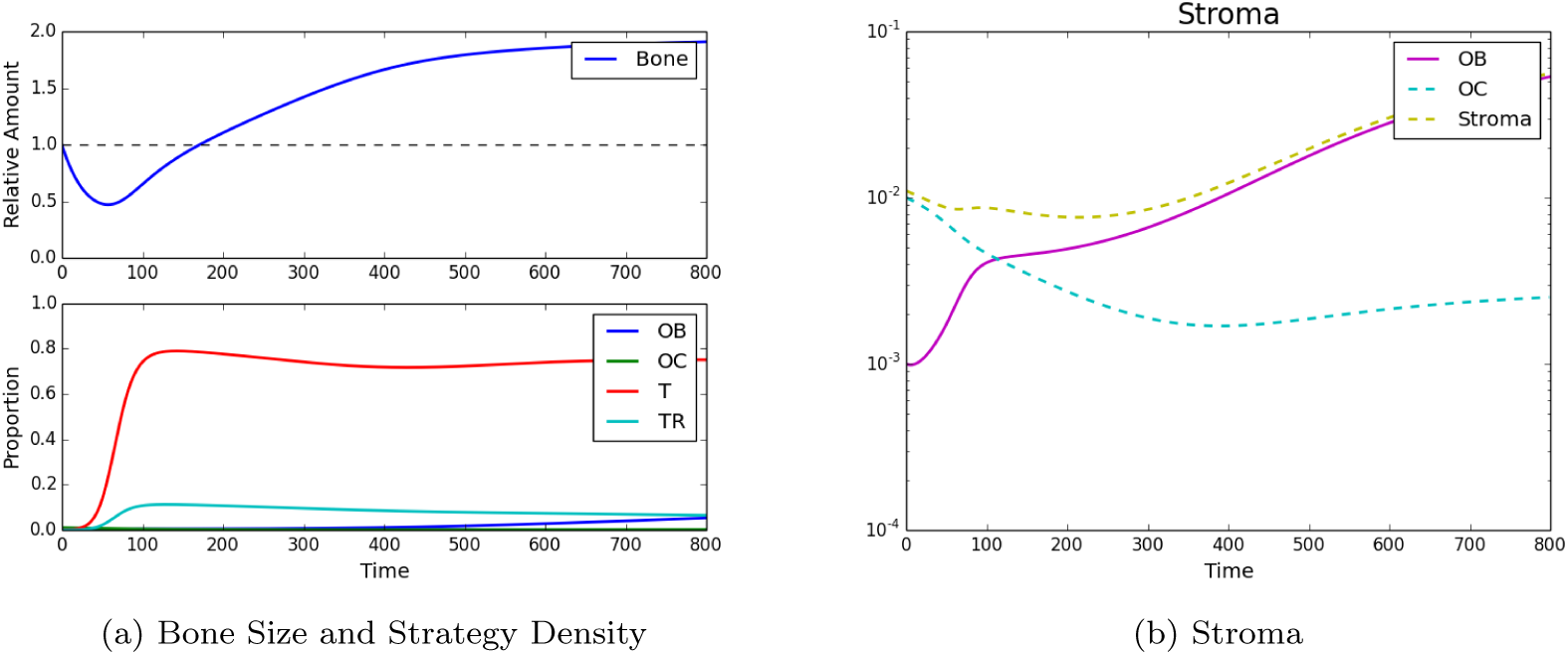
Tumor Introduction. Strategy phenotypes in the bottom of (b) were observed to be dominated by the tumor phenotype. Subfigure (a) and the top of (b) showed the characteristic PCa takeover of the bone remodeling complex and the resultant vicious cycle causing dramatic bone growth.

### Standard Chemotherapy

Trying to asses the impact of chemotherapy we assumed the following initial conditions: *ρ*OB(0) = 0.001, *ρ*OC(0) = 0.01, *ρT* (0) = 0.0005, *ρTR* (0) = 0.0001, *B*(0) = 1. Additionally, a simple chemotherapy regiment of *s* = 0.3 is initiated from timesteps 200 to 350. The results can be seen in figure 3 and show that, while bone continues to grow even under treatment, the tumor population takes a sharp decline highlighting the impact of chemotherapy.

**Figure 3:**
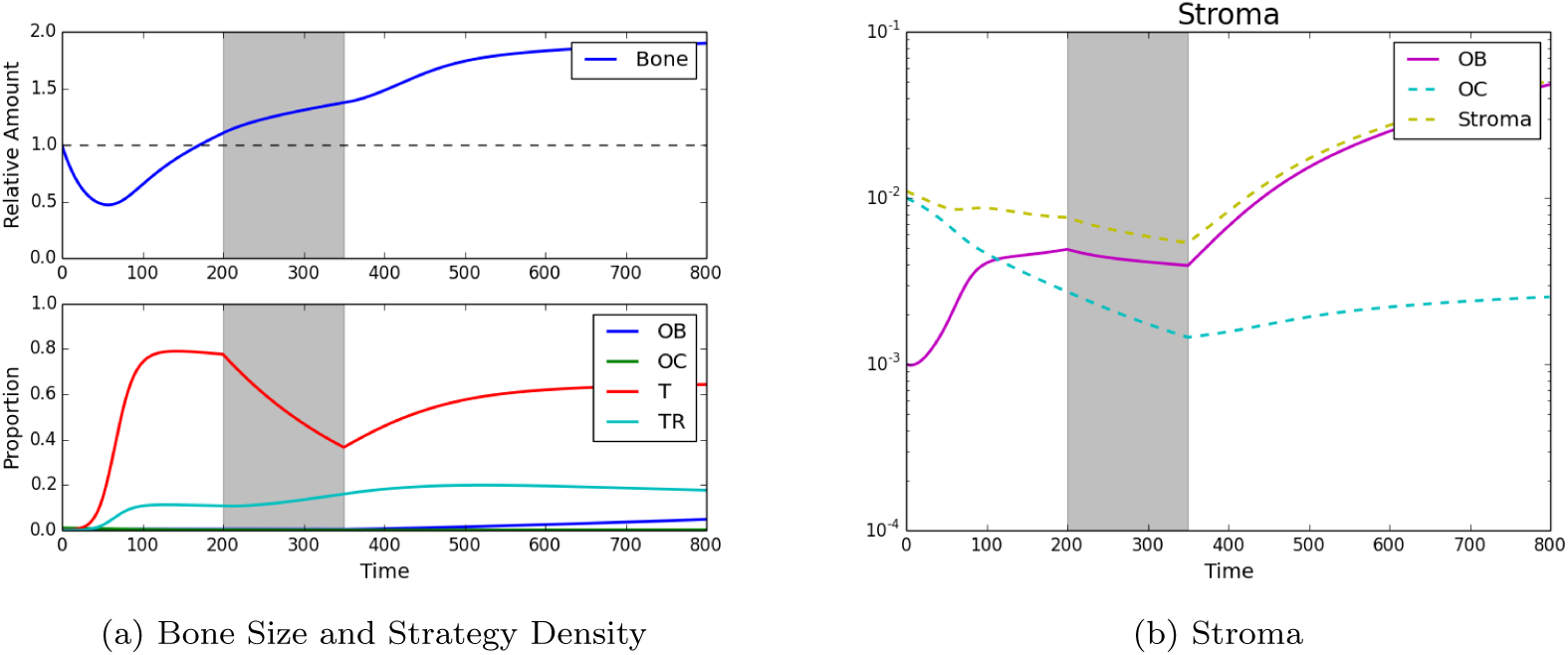
Standard Chemotherapy. Here we see the impact of standard chemotherapy regimens that decrease the PCa susceptible population and increase the resistant PCa population. Assuming initial densities of the different population as follows: *ρ*OB(0) = 0.001, *ρ*OC(0) = 0.01, *ρT* (0) = *ρTR* (0) = 0.0, *B*(0) = 1

### Fractionated Treatment

To understand the effect of varying treatment regimens, the model was used to simulate a wide number of combinations of treatment and treatment holidays. Each regiment is characterized by dividing the treatment into periods, each period representing the same amount of time. Each slot can either have value of (1) to signify that treatment is being applied or (0) when it is not. Thus, assuming a treatment with *n* different treatment periods we can consider 2^*n*^ different treatment regimens based on combinations of treatments and breaks.

The full treatment space also contains regimens that produce favorable results at the cost of eradicating the stromal population - which would be pressumably cytotoxic to the patient. Thus, we removed all treatments that reduced the stromal population below a threshold. This approach allows us to contrast continuous and a spectrum of fractionated regimens. From the space of 16 windows, where each window is 40 timesteps, Table 1 lists the top 5 treatment schedules that yielded the best results in terms of average bone size, as well as the treatment*’*s average tumor burden. Table 1 also lists two other treatment regimens: (1) a continuous treatment regiment and (2) a periodic treatment regiment for comparison.

**Table 1:** Top 5 treatments as well as an example of continuous and periodic treatments. In the 1st column each 0 represents a treatment holiday and each 1 represents the application of chemotherapy. The bone size and tumor burden are dimensionless and rounded to 3 decimal places.

| Treatment Schedule | Bone Size | Average Tumor Burden |
| --- | --- | --- |
| 0000000000000000 | 1.754 | 0.773 |
| 1111100100010010 | 1.160 | 0.352 |
| 1111010101000001 | 1.163 | 0.358 |
| 1111010100100100 | 1.165 | 0.358 |
| 1111011000010010 | 1.166 | 0.356 |
| 1110110101000100 | 1.172 | 0.366 |
| 0001111100000000 | 1.615 | 0.512 |
| 0100100010010011 | 1.709 | 0.666 |

Moreover, figure 4 shows all the potential treatments plotted with respect to average bone size and average tumor burden with three representative treatments shown to the side: a periodic treatment, a continuous treatment and a fractionated treatment.

**Figure 4:**
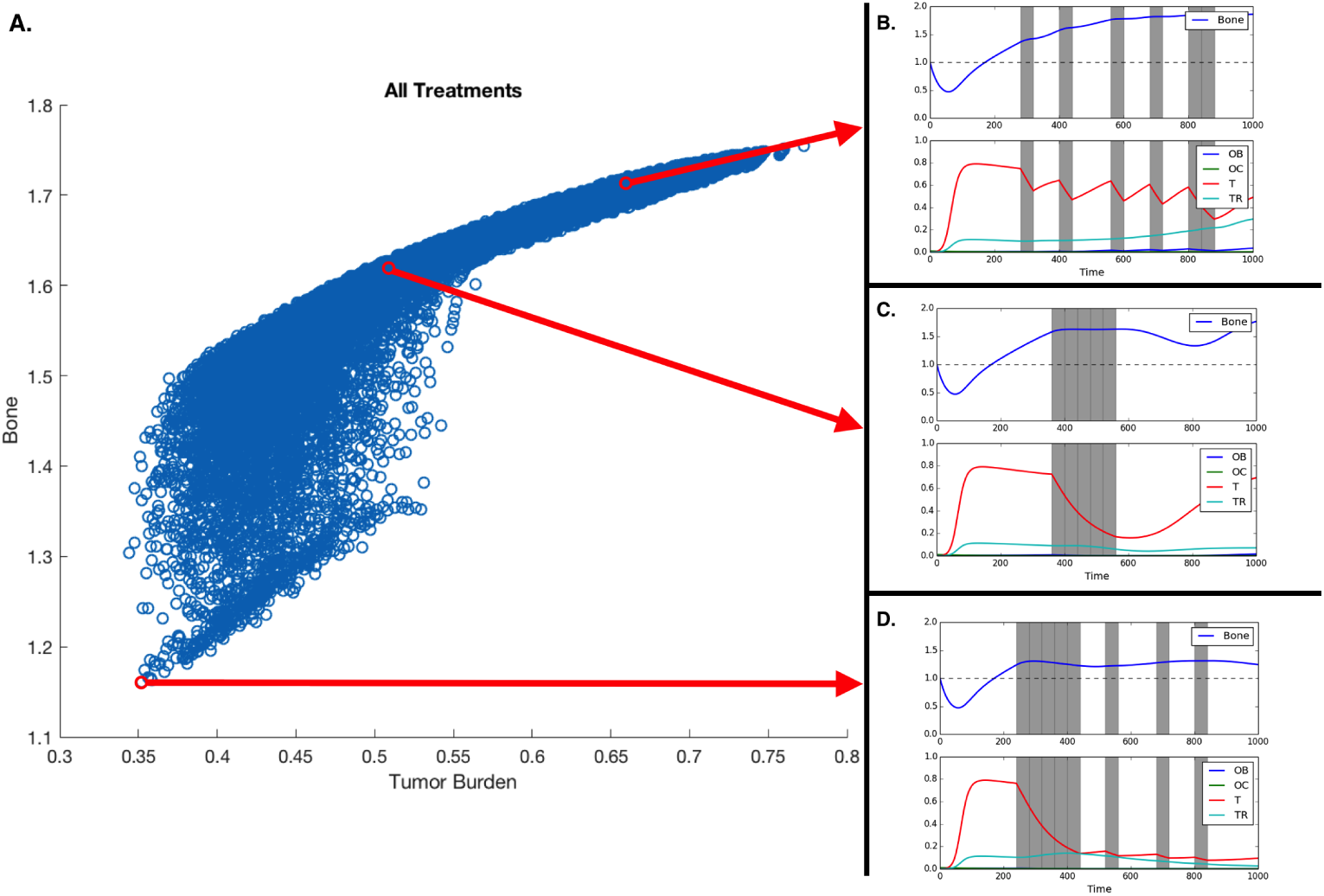
Treatment and treatment holiday combinations. Better treatments are those that reduce tumor burden and extra pathological bone. A. All potential treatments over 16 40-timestep periods, plotted with respect to average tumor burden and average bone size. B,C,D. Treatment regiments showing an example of a fractionated treatment (B), continuous block treatment (C), a fractionated follow-up regiment (D). Assuming initial densities of the different population as follows: *ρ*OB(0) = 0.001, *ρ*OC(0) = 0.01, *ρT* (0) = *ρTR* (0) = 0.0, *B*(0) = 1.

## Discussion

Intra-tumor heterogeneity is increasingly recognized as the key driver of evolution in cancer. This heterogeneity also explains the emergence of resistance to standard of care as well as targeted treatments. We presented an EGT model that captures this heterogeneity and the role of dynamic homeostasis of bone remodeling orchestrateding the interactions between bone-producing cells (osteoblasts) and bone-resorbting cells (osteoclasts). This homeostasis can be disrupted by tumor cells, whose growth, in turn can be prevented by the application of cytotoxic drugs such as docetaxel. Our results recapitulate not only the *”*vicious cycle*”* of the disruption and co-option of homeostatic mechanisms but also the pathological bone formation that characterizes bone metastatic prostate cancer [25]. Knowing that certain tumor phenotypes can be relatively immune to the effects of chemotherapy, our model also captures the emergence of therapy resistant tumors.

The conventional clinical application of chemotherapy in most cancers involves the use of the drug until either the tumor enters remission, resistance renders further application of the drug ineffective or the patient can no longer tolerate it. This approach is called Maximum Tolerable Dosage (MTD) and has been recently contrasted to alternative approaches where the aim is to transform the disease into one that could be managed as a chronic condition. In sipport of this, intermittent docetaxel has been proven to be useful in metastatic prostate cancer [26, 27].

Our results show that the efficacy of the treatment depends on the heterogeneity of the tumor. MTD works best if the metastases are homogeneous. Assuming heterogeneity with regards to chemotherapy resistance, a realistic scenario in metastatic prostate cancer, our model allows us to explore how different treatment duration and intervals between drug application, impact tumor heterogeneity, fitness, and bone mass (see figure 4). Unsurprisingly the model shows that alternative treatment strategies are likely to yield better results if the metastasis contains both susceptible and resistant phenotypes.

Novel therapies where conventional drugs are used while taking into account the tumor*’*s evolutionary dynamics have been proposed and mathematically explored by Orlando et al. [28] under the constraint that the microenvironment of the tumor plays a reduced role and that tumor populations does not interact with each other. Our results extend beyond these constraints and show that neither conventional nor fractionated strategies are always the best solution. While our systematic search has not shown optimal treatments resembling the evolutionary-enlightened therapies proposed by Zhang et al. [22], there is some evidence that they could work in the context of metastatic prostate cancer. Our main result supports a variant: the application of fractionated therapies where full dosage is first applied for a period of time then followed by the alternation between on and off cycles.

While this finding will not have an immediate impact on how fractionated treatments are delivered, it shows that the model captures the basic biological principles of bone metastatic prostate cancer which allowed us to explore treatment strategies in a meaningful way. Models like the one we have presented here can be used to better understand evolutionary-enlightened treatments. EGT is a well known mathematical tool in which to frame evolutionary questions, and our EGT model allows us to include the microenvironmental selection as well as cell-cell interactions in the game dynamics. We are aware that, by its own nature, mathematical models constitute a simplification of the reality being modeled. For example, our model does not take into consideration the spatial interplay which is known to effect EGT dynamics of cancer [14, 29] and has been recently shown to play a key role in the efficacy of some innovative applications of conventional treatments in cancer [30]. We expect that our results could be quantitatively different as we assume different costs of resistance, treatment impact, or spatial structure. But the principle that a period of continuous application followed by periods where on and off cycles is expected to hold when the parameters change to reflect different tumors and treatment efficacies.

We measured the impact of these different treatment schemes by looking at their effect on the two tumor populations but, in bone metastatic prostate cancer, there are other metrics that need to be considered: total dosage and microenvironmental impact. As shown in figure 2, tumor growth leads to an increase in bone, growth that can, in some cases, be curtailed by chemotherapy (see figure 3). Figure 4 shows that the scheduling of chemotherapy can have a substantial impact on the amount of pathological bone formed, a consideration that is clinically important yet rarely included in mathematical models of bone metastases. The examples offer the interesting possibility that a sustained application of chemotherapy followed by treatment holidays and subsequent application of fractionated therapy could lead to control not only on tumor growth but also bone growth (figure 4F; although, see Kaznatcheev [31] for a perspective on the limits of tumour burden control in models like ours). While the total dosage is slightly different, the purely continuous and the (approximately) fractionated treatments shown in figures 4B and 4A demonstrate bone increases over time. Our EGT model, by combining tumor, stroma and bone microenvironment, constitutes a platform in which to investigate various treatment strategies – include, but are not limited, to bisphosphonates, anti-RANKL and hormonal therapies – to the standard of care for metastatic prostate cancer patients that can minimize dosage and bone disease.

## Acknowledgements

We would like to acknowledge Dr. Heiko Enderling from Moffitts Integrated Mathematical Oncology department for the organization of the HIP-IMO program that led to this work. AA, CCL, and DB were partly funded by an NCI U01 (NCI) U01CA202958-01 and a Moffitt Team Science Award. AA was partly funded by a Department of Defense Prostate Cancer Research Program (W81XWH-15-1-0184) fellowship.

